# Subsurface microbial habitats in an extreme desert Mars-analogue environment

**DOI:** 10.1101/269605

**Authors:** Kimberley A. Warren-Rhodes, Kevin C. Lee, Stephen D.J. Archer, Nathalie Cabrol, Linda Ng-Boyle, David Wettergreen, Kris Zacny, Stephen B. Pointing, the NASA Life in the Atacama Project Team

**Affiliations:** NASA Ames Research Center, Moffett Field, CA 94035, USA; The SETI Institute, Mountain View, CA 94043, USA; School of Sciences, Auckland University of Technology, Auckland 1142, New Zealand; College of Engineering, University of Washington, Seattle, WA 98195, USA; Institute of Robotics, Carnegie-Mellon University, Pittsburgh, PA 15213, USA; Honeybee Robotics and Spacecraft Mechanisms Corp., Pasadena, CA 91103, USA; Yale-NUS College and Department of Biological Sciences, National University of Singapore, Singapore 138527

**Keywords:** Atacama, Desert soil, Mars, Moisture stress, Soil bacteria

## Abstract

Sediments in the hyper-arid core of the Atacama Desert are a terrestrial analogue to Mars regolith. Understanding the distribution and drivers of microbial life in the sediment may give critical clues on how to search for biosignatures on Mars. Here, we identify the spatial distribution of highly specialised bacterial communities in previously unexplored depth horizons of subsurface sediments. We deployed an autonomous rover in a mission-relevant Martian drilling scenario with manual sample validation. Subsurface communities were delineated by depth related to sediment moisture. Geochemical analysis indicated soluble salts and minerology that influenced water bio-availability, particularly in deeper sediments. Colonization was also patchy and uncolonized sediment was associated with indicators of extreme osmotic challenge. The study identifies linkage between biocomplexity, moisture and geochemistry in Mars-like sediments at the limit of habitability and demonstrates feasibility of the rover-mounted drill for future Mars sample recovery.

## Introduction

The surface of Mars is dry, cold, and exposed to high levels of ionizing radiation. However, data accumulated over the past few decades by orbital and landed missions have demonstrated that early in its history the planet may have been habitable for microbial life with abundant sources of energy, carbon, nutrients, and shelter (Cabrol, 2018). Mars supported surface and subsurface water and may still do in some circumstances, as well as organic molecules required for life (Martin-Torres et al., 2015). As a result, Mars 2020 and ExoMars missions will be searching for biosignatures (Farley and Williford, 2017; Vago et al., 2017), and the investigation of terrestrial analogues can provide critical insights for the development and testing of exploration strategies. Among those, the hyper-arid core of the Atacama Desert in Chile is widely regarded as a tractable Mars analogue in the field of astrobiology. The Atacama is the driest desert region on Earth (Peel and Finlayson, 2007) with extremely low moisture inputs and precipitation events that are stochastic in nature (McKay et al., 2003). The region has a long history of climatic stability as an extreme desert (Clarke, 2006; Hartley et al., 2005) resulting in the build-up of evaporates (McKay et al., 2003; Navarro-Gonzalez et al., 2003) and creation of a Mars-like surface. The desert’s hyper-arid core lies at or near the arid limit for soil formation (Ewing et al., 2006) and thus surface terrain in this region is regarded as sediment.

Animal and plant life are scarce in extreme deserts and instead cyanobacteria-dominated microbial communities in mineral and rocky refugia including deliquescent substrates comprise the dominant surface biota and are well-characterized (Azua-Bustos et al., 2012; Pointing and Belnap, 2012). Conversely, evidence for microbial colonization in hyper-arid Atacama sediment is scarce, contradictory and almost exclusively limited to surface-associated sediment (Connon et al., 2007; Crits-Christoph et al., 2013; Drees et al., 2006; Lester et al., 2007; Navarro-Gonzalez et al., 2003; Schulze-Makuch et al., 2018) (Supplementary Material, Table S1). Cultivation-based approaches are unreliable as indicators of environmental microbial diversity and have yielded estimates that varied by several orders of magnitude (Bagaley, 2006; Connon et al., 2007; Lester et al., 2007; Navarro-Gonzalez et al., 2003; Schulze-Makuch et al., 2018). Biochemical tests have similarly yielded inconclusive support for microbial metabolic activity (Connon et al., 2007; Crits-Christoph et al., 2013; Lester et al., 2007; Navarro-Gonzalez et al., 2003; Schulze-Makuch et al., 2018). High throughput sequencing of environmental DNA/RNA remains the most reliable indicator for microbial diversity in extreme desert sediments. This approach has provided critical insight on surface-associated communities in semi-arid, arid and hyper-arid locations (Crits-Christoph et al., 2013). A recent study also estimated putative microbial metabolic activity in three subsurface samples after an unusual rain event and this suggested subsurface sediment may also be a habitat for microbial communities (Schulze-Makuch et al., 2018). Major knowledge gaps persist, and importantly from an astrobiology perspective the question of how microbial occurrence and diversity may vary with abiotic variables in sediment depth horizons of a Mars analogue and spatially within terrain, and whether this can be addressed in a simulated robotic sampling mission scenario.

To document these questions and support the development and testing of biosignature exploration strategies, the NASA-funded Subsurface Life in the Atacama (LITA) project deployed an autonomous rover-mounted robotic drill in Mars-like desert pavement and playa terrain within the hyper-arid core of the Atacama under challenging environmental and logistical constraints (Supplementary Material, Fig. S1-S3). The drill accessed 32 discreet sediment samples to depths of 800mm along a 50km transect in a realistic simulation of Martian drilling operations and constraints. Over 60 manually excavated sediment samples were recovered in parallel and validated the automated sampling for abiotic and biotic components of the sediment habitat and provide important baseline ecological data on the most Mars-like sediments on Earth (Warren-Rhodes et al., 2018).

## Materials and Methods

### Field sites

A 50-km autonomous rover traverse in the hyper-arid core of the Atacama Desert was completed in 2013 under challenging environmental and logistical conditions (Supplementary Material Fig. S1). The field experiment took place over a period of two weeks within a landing ellipse circumscribed by rover logistical constraints, ASTER satellite geochemistry and historical climate data as appropriate Mars analogues. The rover traversed two types of terrain identified as appropriate Mars analogues (McKay et al., 2003): stony desert pavement in the western part of the traverse and desert playa in the eastern part within the topographical low and terminus for water runoff (snowmelt, rainfall) for the surrounding region (thus likely to receive significantly more moisture from easterly winter Andean precipitation and runoff than the desert pavement). A unique paleo-playa was also encountered and sampled, comprising an elevated area of dry sediment with evidence of historical playa features. A total of 61 manual samples and 32 rover-acquired samples were independently collected and interrogated.

### Robotic sample recovery

The Zöe rover built by the Robotics Institute at Carnegie Mellon University is a solar-powered rover designed to autonomously map and analyze contextual landscape and habitat visible and geochemical features (with on-board navigation cameras and Vis-NIR spectrometer on its mast) and to drill and deliver samples to on-board scientific instrumentation, including a Mars Micro-beam Raman Spectrometer (MMRS) (Supplementary Material, Fig. S2). The drill, developed by Honeybee Robotics Corporation is a 15kg, 300-Watt, rotary-percussive and fully autonomous drill designed to capture powdered rocks and sediment samples (Online Supplementary Material Fig. S3). The drill consists of a rotary-percussive drill head, sampling augur, brushing station, feed stage and deployment stage, using a vertical 19.1 mm diameter drill operating at 120 rpm. Drilling is accomplished using a bite-sampling approach where samples are captured in 10-20 cm intervals, to simulate a Martian drilling scenario (Zacny et al., 2014). That is, after drilling 10-20 cm, the auger bit with the sample is pulled out of the hole, and the sample brushed off into an on-board sample carousel cup (or sterile Whirlpak (Nasco) bag for manual recovery). Initially autoclaved in the laboratory, the drill bit, brushing station, Z-stage and deployment stage were field sterilized with 70% ethanol prior to and after each site sampling. Aseptic techniques were also used throughout rover sampling operations, including minimal disturbance near the rover during all collections.

### Manual Sediment Pit Sampling

Post-drill, a sediment pit adjacent to the drill hole was excavated manually after first sampling surface sediments. Samples were recovered horizontally from the exposed sediment horizon at 10 cm depth intervals. The pit wall was excavated using a plastic Sterileware (Bel-Art Products) scoop or metal trowel sterilized with 70% ethanol. Deeper layers occasionally required a drill to obtain sample, and a Makita LXT drill was employed, with the drill bit first sterilized with 70% ethanol. All samples were collected using aseptic techniques and tools using a Sterileware (Bel-Art Products) sampling spatula or a stainless-steel spatula sterilized with 70% ethanol. Sediment samples (50-200 g) were collected from each depth layer for biology and geochemistry, with care taken to minimize mixing between different depths, and placed immediately into sterile falcon tubes or Whirlpak (Nasco) bags. Samples were stored at −80°C until processed in the laboratory.

### Mineralogical analysis using rover payload

The MMRS on board the rover collected real-time in-situ mineralogical/geochemical analyses of sediment samples. This allowed direct and unambiguous molecular phase identification from raw Raman spectra of mixtures, e.g., minerals and sediments. The MMRS had a two-unit configuration connected using fiber-optics: A probe unit contained a YAG-KTP crystal pair to generate a 532 nm laser beam for excitation, the optics for laser beam focusing and for Raman photon collection, and a step-motoring mechanism that enable a ∼1 cm line scan at the sample surface. The main unit of the MMRS contained an 808 nm diode pump laser, MMRS Raman spectrometer and detector, electronics for laser driving and control, a microprocessor for commending MMRS operation and data receiving, plus MMRS calibration lamp. This covered a spectral range from 100-4000 cm^-1^ with a spectral resolution of 7–8cm^-1^. The MMRS probe had a working distance of ∼10mm from the sample surface and generated a laser beam of 20–30 mW with a diameter <20 μm at the focus. The wavelength calibration of MMRS spectrometer was performed twice using the Ne calibration lamp during the field campaign, and the laser wavelength calibration was performed daily using naphthalene powder. Raman spectra acquired by the rover were validated against laboratory analysis of sediments using a Kaiser Hololab 5000—532 nm Raman spectrometer with similar configuration to the MMRS and comparable performances in optical throughput (Wei et al., 2015).

### Sediment geochemistry and microclimate data

In the laboratory, sediment chemical analyses for variables known to affect microbial colonization of soil/sediments, including pH, electrical conductivity, soluble salts, C, N, C:N ratio, extractable cations, phosphorous and sulfur plus inherent sediment properties including anion and cation exchange capacity and bulk density were measured according to standard soil chemical analysis methods (https://www.landcareresearch.co.nz/resources/laboratories/environmental-chemistry laboratory/services/soil-testing/methods). It was recognized that high salinity and gypsum levels encountered in our desert sediments may influence estimates for water-soluble ions, however by adopting standard methods applied to soil testing in microbial ecology studies of soil globally we produced a data set that was comparable with other microbial diversity studies.

In brief, geochemical tests were conducted using the following methodology: All sediments were dried in a forced air convection drier at 35°C, and after drying, sediments were crushed to pass through a 2mm sieve. For pH, 10 mL of sediment was slurried with 20 mL of water, and after standing, the pH was measured (1:2 v/v slurry). Bulk density was obtained with a fixed volume (12 mL) of dried and ground sediment weighed. Phosphorus was extracted using Olsen’s procedure (0.5M sodium bicarbonate, pH 8.5, 1:20 v/v sediment:extractant ratio, 30 minutes extraction), and the extracted phosphate was determined calorimetrically by a molybdenum blue procedure. Cations (K^+^, Ca^2+^, Mg^2+^, Na^+^) were extracted using ammonium acetate (1.0M, pH 7, 1:20 v/v sediment:extractant ratio, 30 minutes extraction), and determined by ICP-OES. Cation Exchange Capacity (CEC) was calculated by summation of the extractable cations and the extractable acidity. The extractable acidity was determined from the decrease in pH of the buffered ammonium acetate cation extract. For sulfate-sulfur, sediments were extracted using 0.02M potassium dihydrogen phosphate after 30 minutes shaking, and sulfate-sulfur was measured by anion-exchange chromatography (IC). Total nitrogen (TN) and total carbon (TC) were determined by the Dumas method of combustion. Each sample was combusted to produce varying proportions of CH_4_ and CO gas. The CH_4_ and CO gas was oxidized to CO_2_ using the catalysts Copper Oxide and Platinum. The CO_2_ was measured using Thermal Conductivity detector. Available nitrogen and anaerobically mineralizable nitrogen (not reported) were estimated after incubating sediment samples for seven days at 40°C, after which the ammonium-N was extracted with potassium chloride (2M potassium chloride, 1:5 v/v sediment:extractant ratio, 15 minutes shaking), and determined calorimetrically. A water extraction (1:5 w/v sediment:extractant ratio, 30minutes shaking) was used for electrical conductivity, (EC), which has be used as a proxy for water availability in desert soils/sediments, i.e., higher EC = more salt, less water (Crits-Christoph et al., 2013).

Micro-climate data were collected using *in situ* data loggers in pits installed at the date of sediment sampling in 2013 and recorded continuous subsurface microclimate for 14 months. Relative humidity/temperature HOBO Pro v2 dataloggers (U23-002) were placed at 10 cm, 30 cm and 80 cm depth in pavement and playa sites. Sediment water content was estimated gravimetrically, directly from sediment samples in the laboratory after drying at 120°C x 48hrs to constant dry weight.

### Environmental 16S rRNA gene-defined diversity

Total environmental genomic DNA were extracted from the sediment samples using a method proven during our previous research to maximize recovery from microbial communities from extreme environments. Sediment samples (3x 0.5g triplicate extractions per sample) were combined with 0.5g of 0.1mm silica-zirconia beads. To each sample, 320 µL of phosphate buffer (100 mM NaH_2_PO_4_) and 320 µL of SDS lysis buffer (100 Mm NaCl, 500 mM Tris pH 8.0, 10% SDS) were added and samples were homogenized using the FastPrep^®^-24 (MP Biomedicals, Ohio, USA) at 5.5 for a 30 second run. Samples were centrifuged at 13,200 rpm for 3 minutes (20°C) and 230 µL of cetyltrimethylammonium bromide-polyvinylpyrrolidone (CTAB) extraction buffer (100 Mm Tis-HCl, 1.4 M NaCl, 20 Mm EDTA, 2% CTAB, 1% polyvinylpyrrolidone and 0.4% β-mecaptoethanol) was added. Samples were vortexed for 10 s before incubation at 60°C and 300 rpm for 30 min. Samples were centrifuged at 13,200 rpm for 1 min and then 400 µL of chloroform:isoamyl alcohol (24:1) was added. Samples were then vortexed for 10 s and centrifuged at 13,200 rpm for 5 min. The upper aqueous layer was removed into a new Eppendorf tube and a further 550 µL of chloroform:isoamyl alcohol (24:1) was added. Samples were vortexed for 15 s and again centrifuged at 13,200 rpm for 5 min. The upper aqueous phase was removed into a new Eppendorf tube and 10 M ammonium acetate was added to samples to achieve a final concentration of 2.5 M. Samples were vortexed for 10 s and centrifuged at 13,200 rpm for 5 min. The aqueous layer was removed to a new tube and 0.54 volume of isopropanol was added and mixed by inversion. Samples were left for 24 hours at −20°C and then centrifuged for 20 minutes at 13,200 rpm (4°C). The supernatant was discarded and the pellet was washed with 1 mL 70% ethanol and centrifuged for 10 min at 13,200 rpm. Ethanol was removed and DNA was re-suspended in 20 µL of sterile DNase free water. Samples were then quantified using the Qubit 2.0 Flourometer (Invitrogen). Samples were then stored at −20°C until required. The extracted DNA were then adjusted, where possible, to 5 ng/μL before Illumina MiSeq library preparation as specified by the manufacturer (16S Metagenomic Sequencing Library Preparation Part # 15044223 Rev. B; Illumina, San Diego, CA, USA). Briefly, PCR was conducted with the primer set targeting the V3-V4 regions of bacterial and archaeal 16S rRNA gene: PCR1 forward (5′ TCGTCGGCAG CGTCAGATGT GTATAAGAGA CAGCCTACGG GNGGCWGCAG 3′) and PCR1 reverse (5′ GTCTCGTGGG CTCGGAGATG TGTATAAGAG ACAGGACTAC HVGGGTATCT AATCC 3′) with KAPA HiFi Hotstart Readymix (Kapa Biosystems, Wilmington, MA, USA) and the following thermocycling parameters: (1) 95°C for 3 min, (2) 25 cycles of 95°C for 30 s, 55°C for 30 s, 72°C for 30 s, 72°C for 5 min, and (3) holding the samples at 4°C. The amplicons were then indexed using Nextera XT index kit (Illumina). The indexed amplicons were purified and size selected using AMPure XP beads (Beckman-Coulter, Brea, CA, USA) before sequencing on an Illumina Miseq (Illumina) with the 500 cycle V2 chemistry (250 bp paired-end reads). A 5% PhiX spike-in was used, as per manufacturer’s recommendation. The R packages phyloseq, DESeq2 and ggplot2 were used for downstream analysis and visualization including ordination and alpha diversity calculations (https://www.r-project.org/). High-throughput sequencing of the 16S rRNA gene yielded 87,8875 quality filtered reads and 92 bacterial OTUs that were further analyzed. All sequence data acquired during this investigation has been deposited in the NCBI Sequence Read Archive under project accession number PRJEB22902.

Our approach employed an environmental DNA recovery method that has been optimized for extreme desert sediments and has been used as a proxy for relative biomass(Pointing et al., 2009). We employed positive and negative controls for all PCR amplifications. We employed rRNA gene loci widely accepted as the benchmark for interrogating environmental soil/sediment samples (http://www.earthmicrobiome.org/). We also adopted high coverage and carefully screened our sequence libraries for artefacts and contaminants. We also performed successful DNA extractions on pavement and playa sediments that yielded negative extraction outcomes for environmental DNA (as well as those yielding positive outcomes as controls), after spiking these with *E.coli* from an axenic cell suspension in phosphate buffered saline solution at final cell concentrations of 10^3^ – 10^7^ cfu /g sediment to simulate estimated occurrence in hyper arid sediment (Connon et al., 2007; Lester et al., 2007). Recovery rates were in the range of 60-80% of that for axenic cell suspensions in all treatments.

We did not supplement our intensive molecular ecological survey with a cultivation approach because it is widely accepted that most environmental bacteria are unculturable and their *in vitro* growth requirements are unknown, thus such an approach would have introduced unacceptable bias to the study. Similarly, the lack of resolution and triangulation available for some alternative approaches including lipid analysis and metabolic activity limit their contribution to microbial community estimation relative to DNA sequencing approaches. Given the inherent patchiness of distribution for microbial life in extreme deserts and our robust methodological approach we are confident that our estimates provided the most accurate possible indication of endemic diversity.

***Statistical analysis***

Canonical correspondence analysis (CCA) was performed with the R package vegan to explore the strength of associations among the sediment geochemistry profiles, bacterial taxa (OTUs) and site locations (https://www.r-project.org/). Type III symmetrical scaling was used in the CCA plot, where both site and species scores were scaled symmetrically by square root of eigenvalues. This technique provided a weighted sum of the variables that maximizes the correlation between the canonical variates. A biplot was created to visualize the outcomes and help facilitate interpretation of the canonical variate scores. Linear Discriminant Analysis (LDA) using the R package flipMultivariates (https://github.com/Displayr/flipMultivariates) was applied to geochemistry data (Online Supplementary Table S3) (n=61, cases used for estimation n=47, values below detection range were treated as 0, cases containing missing values were excluded, null hypotheses = two-sided). For multiple comparisons correction, False Discovery Rate correction was applied simultaneously to the entire table. BEST analyses were conducted using the BIO-ENV procedure in Primer 7 software (http://www.primer-e.com/) to maximize the rank correlation between biotic and environmental data, thereby establishing a ranking (_*p*_*w*) for the effects of environmental variables on diversity.

## Results and Discussion

### Extreme habitats in Atacama sediment horizons

The on-board Raman spectrometry revealed surface mineral distributions comprising feldspar and quartz. Subsurface sediments, and particularly the playa sites, were elevated in gypsum and anhydrite (Supplementary Material, Table S3, Fig. S5). This was independently corroborated by laboratory Raman measurements (Supplementary Material, Table S3) and geochemical analysis that revealed a positive linear correlation (r=0.78, p<0.001) across all samples between sulfate-sulfur and calcium, the components of gypsum and anhydrite (CaSO_4_ +/-2H_2_O). Anydrite is indicative of drier conditions at depth, and *in situ* soil moisture sensors validated this for horizons in both pavement and playa.

Our chemical analysis focused on geochemical reservoirs readily available and relevant to biological communities and using methodology that was comparable with other microbial ecology studies. A clear depth-dependent pattern in geochemistry was revealed for pavement and playa horizons (Fig. 1, Supplementary Material, Table S2, Fig. S4). Surface sediments were strongly associated with elevated phosphorous concentrations in both playa and pavement units (Fig. 1). Elevated surface P was attributed to presence of surface Fe-oxides and calcite that can adsorb P. These P-adsorbing phases were likely present in the subsurface but were diluted by high sulfate concentrations causing lower recorded P concentrations below the surface. All subsurface samples displayed very low to undetectable levels of N and low C throughout horizons. The on-board Raman spectrometer also recorded C as a minor/undetectable phase in all samples. Electrical conductivity (EC) and Na^+^ levels increased with depth in both substrates indicating increased salinity with depth (p = 0.006). Desert pavement subsurface sediments separated mainly due to pH and K^+^, whilst playa subsurface sediments displayed elevated levels of sulfate-sulfur, extractable cations (Ca^2+^, Mg^2+^, Na^+^) and EC indicating increasing osmotic challenge, and most notably in the paleo-playa Site 10 (Fig. 1, Supplementary Material Table S2).

**Figure 1.**
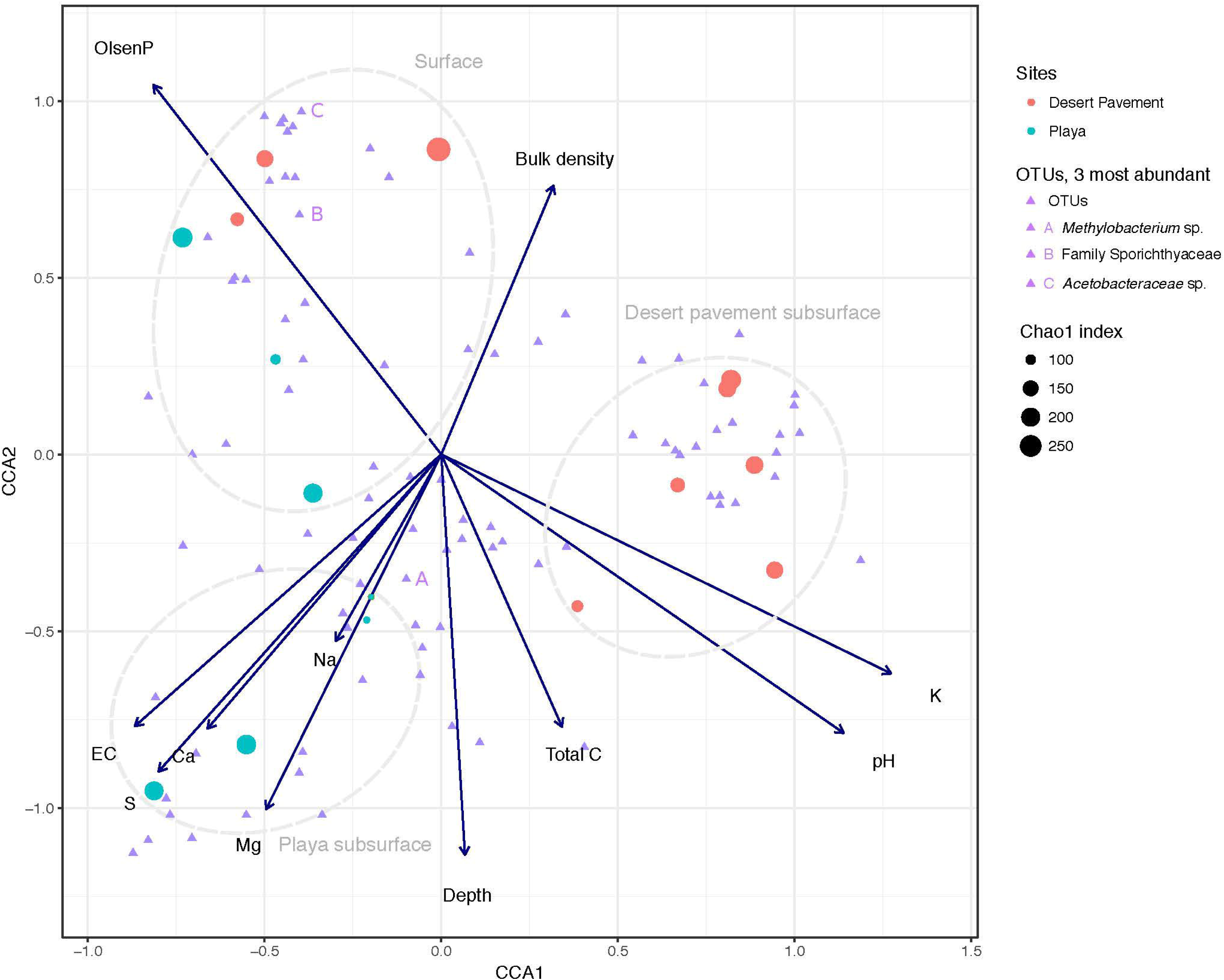
Correlation of Mars analogue sediment geochemistry with bacterial diversity. Canonical Correspondence Analysis (CCA) triplot with symmetrical scaling indicating differences in sediment geochemistry within sediment pits, and influence of these abiotic variables on bacterial communities and individual taxa. The three most abundant taxa are labelled (A, B, C). The circle size of each sample indicates species richness index (Chao 1) of the respective community.

Average annual subsurface temperatures ranged from 19.9 – 20.9°C in depth horizons (Supplementary Material, Table S4). At shallow depths (<100mm) temperature varied between 4 – 34°C, whereas in deeper horizons temperature variation was somewhat buffered (Supplementary Material, Fig. S6). Moisture values from depth horizons in representative playa terrain were consistently higher (approximately 4-to 7-fold) at all depths than those in the desert pavement horizons (Supplementary Material, Table S4) and this likely reflects local hydrology given the playa was a relatively low-lying terrain (Supplementary Material, Fig. S1). However, sediment moisture trends in both substrates broadly indicated the existence of depth groupings into distinct moisture zones: (a) a surface zone consisting of the top 200mm, where water availability is typically lowest, except in the short-term following a rain event; (b) a mid-depth zone (300-500mm), where water availability peaks and persists after a rain event; and (c) a deep subsurface zone (≥500-800mm), where water availability is typically lower and, notably for the desert pavement sediments, appeared to be un-impacted by rare large rainfall events (Supplementary Material, Table S4, Fig. S6).

### Depth-related trends in DNA recovery and microbial alpha diversity

Surface colonization was widespread in desert pavement and playa sites but subsurface horizons displayed patchy recovery with low yields of quantifiable DNA in the range 0.067-6.5 ng/g sediment, indicating extremely low standing biomass (Supplementary Material, Table S2, S5). Notably, the paleo-playa site yielded no recoverable DNA at any depth after repeated efforts at extraction. Our approach using a DNA recovery method adapted for low-biomass extreme environments highlights for the first time the patchiness of low biomass microbial colonization in the most extreme subsurface desert sediments where micro-habitat conditions are at or near the limit for life (McKay, 2014). This may explain, at least in part, why some previous research has concluded this region of the Atacama was lifeless (Navarro-Gonzalez et al., 2003). Linear Discriminant Analysis was employed to show that variables most strongly associated with potentially lifeless subsurface sediments where environmental DNA was irrecoverable were sulfate-sulfur (substrate) (p = 0.001), depth (p = 0.003), EC (p = 0.006), soluble salts (p = 0.006). This strongly suggests that osmotic challenge and limited moisture availability are the major extinction drivers in this Mars analogue sediment. We also demonstrated that substrate-inhibition of DNA recovery was unlikely to have been a significant factor, since bacterial cell suspensions in the range reported for desert soils (Supplementary Material, Table S1) added to these samples were recoverable with 60-80% efficiency compared to a pure laboratory culture and were not significantly inhibited compared to extractions with spiked sediments that did originally yield environmental DNA.

Across both terrains any given depth horizon tended to either be colonized almost throughout or display only patchy near-surface colonization as measured by recoverable DNA. Thus, the ubiquitous surface colonization was not a predictor for subsurface habitability in these sediments. We postulate that in addition to sediment moisture, extreme geochemistry also influenced habitability in these sediments as evidenced by high soluble salts/salinity indicators and abundance of anhydrite and gypsum mineral signatures (Supplementary Material, Table S2, S3). Temperature variations were not regarded as significant challenges to microbial colonization compared to other variables (Supplementary Material, Fig. S6). The paleo-playa site 10 failed to yield recoverable DNA from any depth, thus highlighting that ancient landforms created under moisture sufficiency, and with elevated anhydrite levels as indicated by the rover’s Raman spectrometer (Supplementary Material, Fig. S5) may not support extant genetic biosignatures and do not support recoverable relic DNA. Our overall DNA recovery success was consistent with our expectations for the driest desert location on Earth. It is notable that we have observed Antarctic mineral sediments, another Mars analogue, generally yield higher recovery rates using similar methodologies, e.g. (Lee et al., 2012; Pointing et al., 2009). However, this is due to enhanced standing biomass due to the less extreme nature of the growing season in Antarctic desert where long periods of frozen hibernation are punctuated by periods of moisture sufficiency during the austral summer (Pointing et al., 2015).

Clear patterns were discernible from manually excavated sediment pits. Bacterial diversity decreased significantly with depth across both terrains (Chao1 richness: r = 0.508, p = 0.044; Shannon’s H: r = 0.731, p = 0.001), thus indicating depth as a major driver of bacterial diversity (Supplementary Material, Fig. S8, Table S6). Bacterial communities formed six diversity clusters that associated with clearly defined depth ranges and zones of variability in sediment moisture and geochemistry (Fig. 2a). Subsurface communities were more distinct between habitat types but also more heterogenous overall than surface-associated communities, due largely to the lower bacterial diversity within each depth-defined sediment micro-habitat. Community evenness displayed little clear pattern with depth for desert pavement but for playa deep samples displayed very low evenness reflecting their highly specialized diversity (Supplementary Material, Table S6). In all cases the rover-acquired samples showed a generally similar pattern in depth profile and diversity clusters although these were less clearly defined and we attributed this to mixing of sediment during recovery using the bite-drilling approach (Fig. 2b; Supplementary Material, Fig. S8).

**Figure 2.**
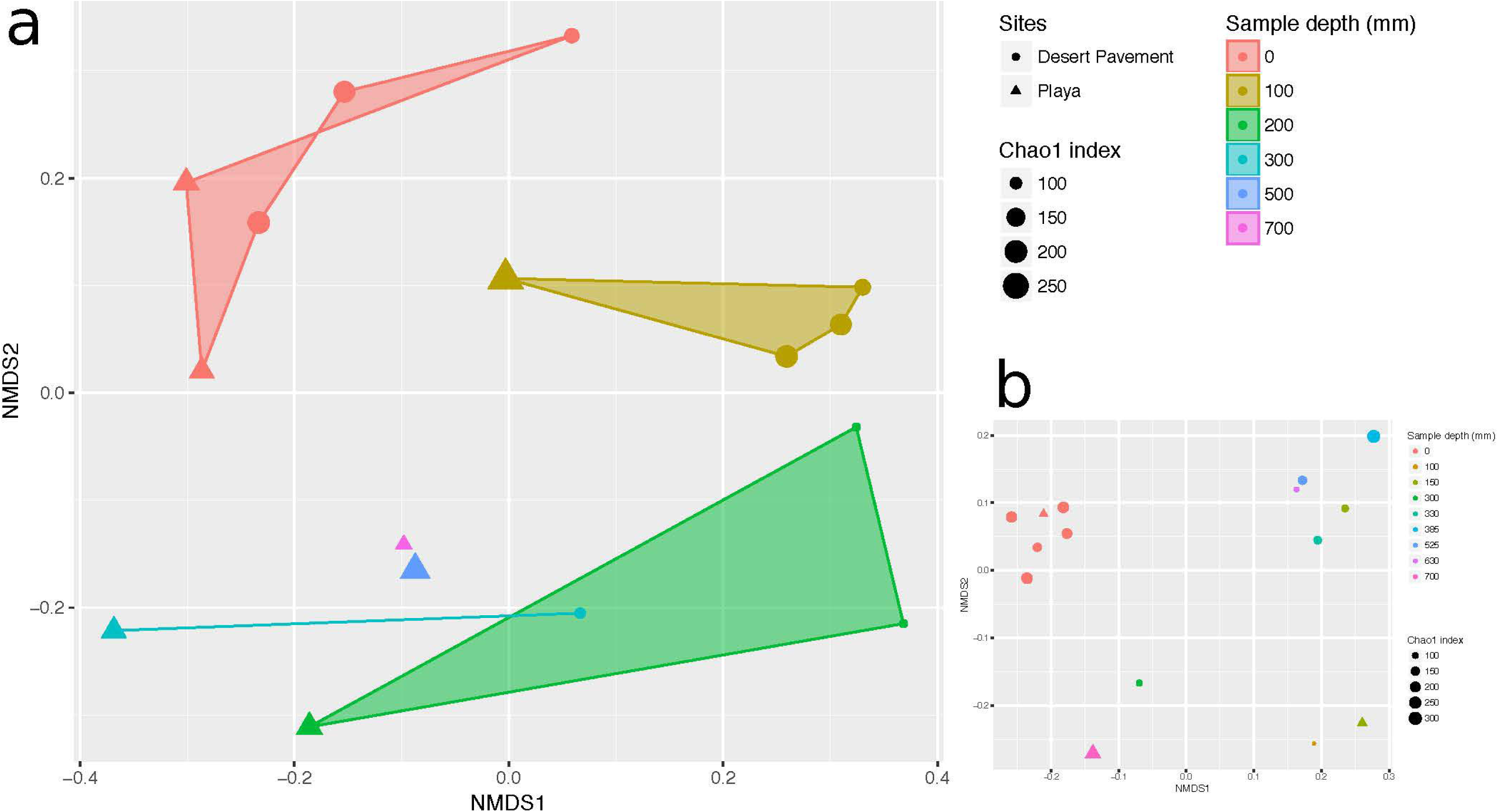
Depth-defined bacterial diversity in Mars analogue sediments of the Atacama Desert. Non-metric multidimensional scaling (NMDS) ordination of Bray Curtis similarities for bacterial diversity versus sediment depth from a) manual recovery, and b) rover recovery. Shaded areas indicate similarity clusters for sediment communities at the same depth. The size of each symbol (circle or triangle) indicates species richness index (Chao 1) of the respective community. Mid-range values were used for drill samples where depth ranges instead of individual depths were generated.

To further unravel the influence of sediment environment on bacterial diversity, we performed canonical correspondence analysis (CCA) to establish the association of distinct geochemistry for desert pavement and playa samples on the assembly of bacterial communities (Fig. 1). Surface communities were strongly influenced by P levels that were elevated compared to the subsurface (Fig. 1). Bacterial diversity in subsurface sediment habitats correlated with two groups of geochemical variables: The playa subsurface community was strongly associated with EC and extractable cations (Ca^2+^, Mg^2+^, Na^+^) and the rover-mounted Raman data indicated elevated anhydrite and gypsum (Supplementary Material, Table S3, Fig. S5). This strongly suggests that bio-availability of water may be limited in this subsurface habitat despite relatively high moisture levels due to the effects of salt saturation. Conversely the pavement subsurface community was associated largely with pH, and also C and K^+^ although the magnitude of variation for these variables was low. This pH-dependent structuring of microbial diversity is consistent with observations for global trends in soil microbial diversity (Fierer and Jackson, 2006). The BEST multiple rank correlation routine was employed to further examine our data and rank the relative association of abiotic variables with the observed bacterial diversity as follows: extractable cations (Ca2+, Mg2+, Na+) and phosphorous>sulfate-sulfur> >EC (_*p*_*w* = 0.595-0.609, *p* <0.05), thus further validating the ordinations and CCA analysis.

### Highly specialized sediment bacterial communities

The taxa identified in sediment indicated highly specialized and relatively low diversity bacterial communities in both desert pavement and playa, and this low diversity reflects global trends in soil microbial diversity where deserts are considered to be relatively depauperate (Delgado-Baquerizo et al., 2018; Thompson et al., 2017). Overall the drill samples yielded weaker depth resolution than manual sampling but still corroborated observations from the manually collected samples (Fig. 3, Supplementary Material Fig. S9). Bacterial taxonomic diversity varied significantly with subsurface habitat as compared to a generally consistent surface community (Fig. 3) (Crits-Christoph et al., 2013). Overall, communities were dominated largely by only three phyla: Chloroflexi, Actinobacteria and Alphaproteobacteria.

**Figure 3.**
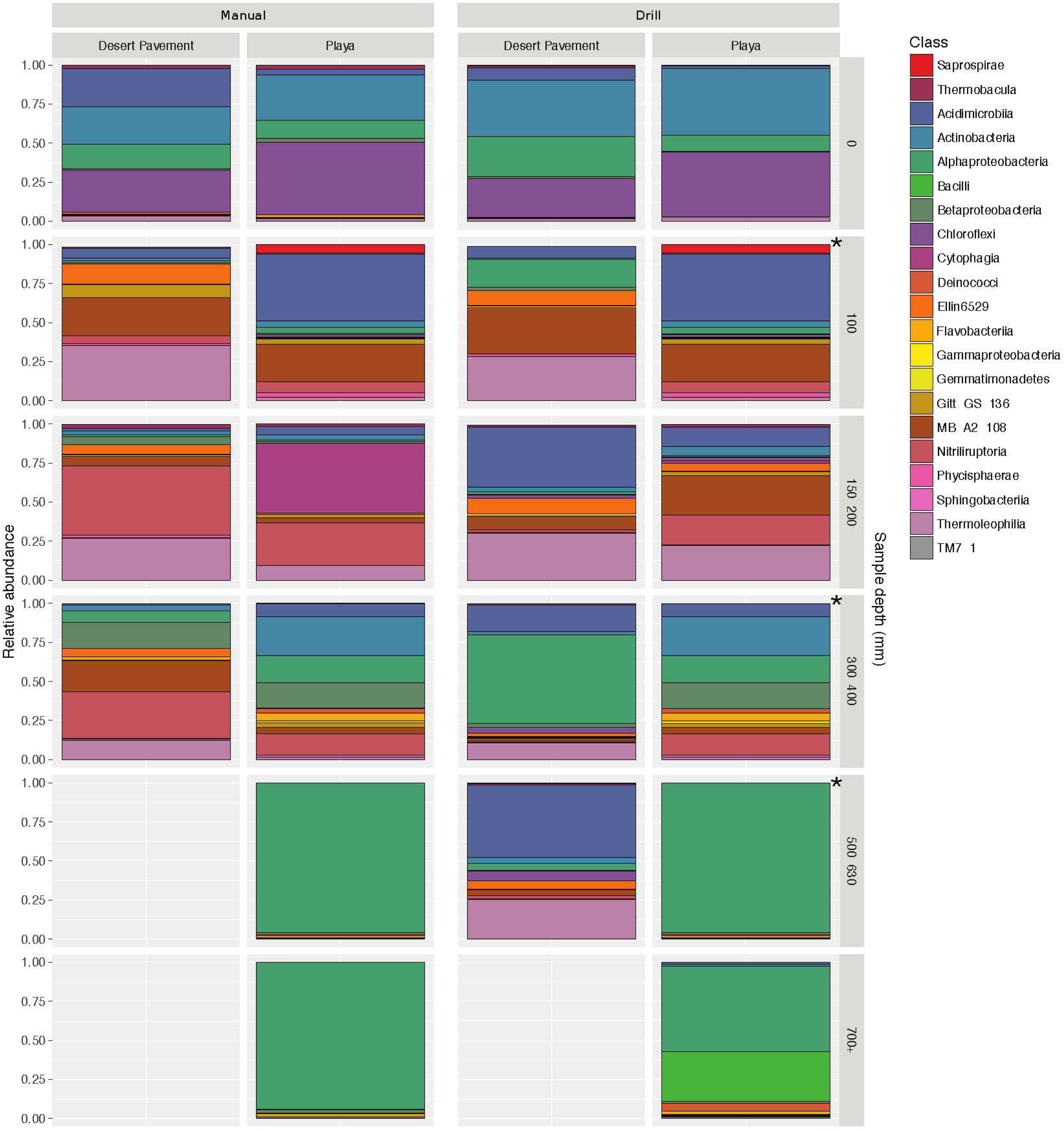
Highly specialized bacterial communities in Mars analogue sediment depth horizons. Distribution of bacterial diversity by taxonomic class with sediment depth for manual (M) and autonomous rover drill (D) recovered samples. Colored shading indicates relative abundance within each community for a given bacterial class. Grey shading indicates no recoverable bacteria.

All surface communities were dominated by the AKIW781 lineage of the photoheterotrophic Chloroflexi, a known desert soil crust taxon (NCBI GenBank accession number: JQ402146), and overall community structure contributed additional taxonomic resolution to previous observations of surface sediment communities (Crits-Christoph et al., 2013; Schulze-Makuch et al., 2018). AKIW781 has also been recorded in desert varnish on rock surfaces (Kuhlman et al., 2008) as well as a keystone taxon of hypolithic communities (Lacap et al., 2011) in the Atacama. This indicates a cosmopolitan distribution and broad habitat preference among surface niches in this extreme desert. Conversely, the Chloroflexi were minor components of subsurface communities, decreasing in relative abundance with depth, and mainly comprised an uncharacterized candidate class Ellin6529, likely adapted to non-phototrophic metabolism in the subsurface microhabitat.

Low and mid-depth subsurface horizons for both desert pavement and playa were dominated by the low G-C gram positive Actinobacteria (Fig. 3, Supplementary Material, Fig. S9). They are typically thick-walled bacteria commonly encountered in soil. Communities encountered were specific to each depth horizon and shifts in diversity clearly reflected the moisture zones identified for both pavement and playa horizons. At shallow depths (<200mm) halotolerant, alkalotolerant, spore-forming desiccation-tolerant actinobacterial groups were abundant and included the orders acidimicrobiales, gaiellales, nitriliruptorales and solirubrobactales in desert pavement and acidimicrobiales and nitriliruptorales in playa. Mid-depth sediments (300-<500 mm) corresponding to the zone of greatest moisture availability in both substrates supported highly variable bacterial diversity between samples and also the most diverse communities (Supplementary Material, Table S6). Of note was the elevated relative abundance of bacillales at a single pavement site and this may reflect the relatively high altitude and limited moisture input to the subsurface. The bacillales-dominated community was more similar to those encountered at greater depths at other locations. The bacillales are characterized by their ability to form highly resistant endospores and so may indicate an increasingly challenging micro-habitat in deeper subsurface sediments.

There were fewer deep sediment samples from which to make comparisons and this reflected the more challenging subsurface environment, and deeper communities (500+ mm) generally displayed lowest taxonomic diversity. The deepest desert pavement community was similar to those at shallower depths but displayed elevated relative abundance of acidimicrobiales. Conversely the deep playa community shifted to a fundamentally different composition from shallower depths that was dominated almost exclusively by a single facultative methylotrophic and desiccation-tolerant *Methylobacterium radiotolerans* taxon (NCBI GenBank accession number: LT985974) from the phylum Alphaproteobacteria. We speculate the C1 metabolism of this taxon allows it to exploit simple C1 compounds as well as subsurface methane sources (Kao et al., 2017), a molecule known to be released from subsurface sources on Mars (Stevens et al., 2017). An overall picture thus emerges of highly-specialized bacterial diversity adapted to and reflecting the challenging subsurface habitat as well as reflecting moisture availability zones determined at least in part by sediment depth and geochemistry.

Other bacteria typically encountered in desert surface communities and generally regarded as tolerant to extreme conditions were not major components of subsurface communities. A complete absence of cyanobacterial taxa was consistent with their adaptation to surface mineral refugia rather than subsurface sediment habitats that preclude photoautotrophy (Azua-Bustos et al., 2012), and the highly desiccation-tolerant *Deinococcus*-Thermus group were represented by only a single lineage of Trueperaceae candidate genus B-42 recovered in just a few subsurface samples with low abundance (0.6 – 4.8%), compared to a relatively diverse assemblage of cultivable Deinococci recovered from surface sediments in other less arid desert locations (Rainey et al., 2005). This further supports our evidence for highly specialized subsurface communities selected for by the distinct geochemistry and microclimate in the subsurface Mars analogue sediments of the Atacama. They may thus be broadly indicative of the type of microbial consortia that could exploit Martian subsurface habitats.

In the deepest and least-diverse playa communities biocomplexity was reduced almost to a single taxon, reflecting extreme selective pressure and also highlighting a possible lack of resilience to environmental change given that recruitment to deep subsurface sediments may be limited. The minimum biocomplexity may comprise multiple ecotypes of a single taxon, each adapted to exploit a given suite of microclimate and geochemical conditions (Koeppel et al., 2008). They exhibited a strong preference for C1 and/or autotrophic taxa that are somewhat delinked from their immediate surroundings in terms of carbon sequestration and reflect the extreme oligotrophic nature of these microhabitats. Given the highly specialized diversity recovered, the association of putative physiology with environmental variables, and lack of DNA recovery in paleo-playa samples; we conclude that our estimates are likely to represent resident microbial communities rather than relic DNA (Carini et al., 2016; Schulze-Makuch et al., 2018).

### Implications for detection of biosignatures on Mars

The autonomous rover drilling platform yielded subsurface sediment samples that allowed for the first time a combined investigation of sediment abiotic properties and microbial diversity at an unprecedented level of detail. The parallel manual sampling and analysis demonstrated feasibility and fidelity of the autonomous rover approach. Highly specialized but low-diversity subsurface bacterial communities were encountered patchily and this was strongly associated with abiotic variables which suggests that “follow the water” is only part of the biosignature exploration solution in the search for potential habitable refuges on Mars. Consideration of subsurface micro-habitat variability in geochemistry, originating with and adapted to possible water availability zones may also be key. Whilst the geochemistry of our analogue sites was similar to that of a habitable Martian regolith, moisture in the Atacama is surface-sourced by fog and/or rain events (McKay et al., 2003), whereas on Mars subsurface sources may provide an upward migration of moisture similar to that observed in Antarctic mineral sediment overlaying permafrost (Goordial et al., 2016; Stomeo et al., 2012). Thus, extrapolating habitable subsurface locations on Mars would need to consider this along with the incident radiation regime and other Martian environmental variables. Based on the exploratory design consideration for this NASA-funded research, drill depth was constrained at 800mm as a proof of concept study. However, current plans are for both NASA and ESA to target depths of up to 2m on Mars in order to target potential subsurface habitats that account for Martian environmental variables.

The relevance of ecology and microbial habitats to past and possible extant life on Mars are finally coming to the fore in the robotic search for biosignatures on Mars (Cabrol, 2018; Warren-Rhodes et al., 2007). As our study suggests, detecting such life or its residual biosignatures may prove highly challenging, given that in the most extreme deserts on Earth these communities are extremely patchy in distribution and occur with exceedingly low biomass. The drill apparatus employed in this study has demonstrated that subsurface sediment biosignatures can be autonomously recovered, although precise depth delineation requires refinement with the bite-drilling approach used in this study. Whilst genetic biosignatures such as DNA used in our study may not ultimately be the primary method employed to search for traces of life on Mars, they are the most reliable and widely used method currently available for microbial diversity estimation (Delgado-Baquerizo et al., 2018; Thompson et al., 2017). This approach provided essential first proof of concept that an incontrovertible biological signature within the likely range for geochemical variables in a habitable subsurface environment can be recovered from a Mars-like sediment using an autonomous rover.

## Acknowledgements

The work was completed with funding from NASA Astrobiology Science and Technology for Exploring Planets (ASTEP) Program Grant NNX11AJ87G and a Yale-NUS College Start-Up Grant. NASA Life in the Atacama Project team members that contributed to the success of this research: Guillermo Chong (Universidad Católica del Norte), Cecilia Demargasso (Universidad Católica del Norte), Greydon Foil (Carnegie-Mellon University), Christopher Gayle Tate (University of Tenessee), Trent Hare (US Geological Survey), Donnabella C. Lacap-Bugler (Auckland University of Technology), Jeff Moersch (University of Tenessee), Ken Tanaka (US Geological Survey), Cinthya Tebes (Universidad Católica del Norte), Srinivasan Vijayarangan (Honeybee Robotics and Spacecraft Mechanisms Corp.), Michael Wagner (Carnegie-Mellon University), Alian Wang (Washington University), Jie Wei (Washington University). The authors are grateful to Brad Sutter (NASA) for advice on interpretation of geochemical analysis, and Craig Cary (University of Waikato) for assistance with biosecurity compliance and sample custody. Author Kris Zacny was employed by Honeybee Robotics and Spacecraft Corporation. All other authors declare no competing interests. A preprint version of this manuscript is available on the bioRxiv server: https://www.biorxiv.org/content/early/2018/06/20/269605.

